# Electrophysiological correlates of mood and reward dynamics in human adolescents

**DOI:** 10.1101/2021.03.04.433969

**Authors:** Lucrezia Liuzzi, Katharine K. Chang, Hanna Keren, Charles Zheng, Dipta Saha, Dylan M. Nielson, Argyris Stringaris

## Abstract

Despite its omnipresence in everyday interactions and its importance for mental health, mood and its neuronal underpinnings are poorly understood. Computational models can help identify parameters affecting self-reported mood during mood induction tasks. Here we test if computationally modelled dynamics of self-reported mood during monetary gambling can be used to identify trial-by-trial variations in neuronal activity. To this end, we shifted mood in healthy (N=24) and depressed (N=30) adolescents by delivering individually tailored reward prediction errors whilst recording magnetoencephalography (MEG) data. Following a pre-registered analysis, we hypothesize that expectation (defined by previous reward outcomes) would be predictive of beta-gamma oscillatory power (25-40Hz), a frequency shown to modulate to reward feedback. We also hypothesize that trial variations in the evoked response to the presentation of gambling options and in source localized responses to reward feedback. Through our multilevel statistical analysis, we found confirmatory evidence that beta-gamma power is positively related to reward expectation during mood shifts, with possible localized sources in the posterior cingulate cortex. We also confirmed reward prediction error to be predictive of trial-level variations in the response of the paracentral lobule and expectation to have an effect on the cerebellum after presentation of gambling options. To our knowledge, this is the first study to relate fluctuations in mood on a minute timescale to variations in neural oscillations with noninvasive electrophysiology.

**Significance Statement:** Brain mechanisms underlying mood and its relationship with changes in reward contingencies in the environment are still elusive but could have a strong impact on our understanding and treatment of debilitating mood disorders. Building on a previously proposed computational mood model we use multilevel statistical models to find relationship between trial-by-trial variations in model components of mood and neural responses to rewards measured with non-invasive electrophysiology (MEG). Through confirmatory analysis we show that it is possible to observe relationships between trial variations in neural responses and computational parameters describing mood dynamics. Identifying the dynamics of mood and the neural processes it affects could pave the way for more effective neuromodulation treatments.

## Introduction

Humans report on their moods in their everyday conversations and subjective mood reports form the basis of clinical assessment and much of research in affective neuroscience. Advances in computational modeling lend support to the idea that mood is intricately linked with reward processing and that it serves to integrates over a person’s history of rewards and punishers (Keren et al., 2020; Nettle and Bateson, 2012; Rutledge et al., 2014). Yet, despite its ubiquity and importance, the brain mechanisms underlying mood and its relationship with changes in reward contingencies in the environment are surprisingly understudied.

Mood is understood to integrate over events in the environment and is a potentially emergent property of the coordinated activity of many neural populations. Such activity is thought to manifest as the synchrony of oscillations which supports functional connections and communication in the brain. In this context, it is noteworthy that oscillatory power is correlated with treatment-induced changes that have been described to occur in mood disorders (Fingelkurts and Fingelkurts, 2015; Kaiser et al., 2015; Nugent et al., 2019a, 2019b). Mood is also highly dynamic during development, especially in adolescence (Klimstra et al., 2016). Understanding the temporal structure—including very early responses—to changes in environmental incentives that influence mood can offer important insights on the genesis and remission of mood disorders and can inform the timing of potential interventions (such as through transcranial magnetic stimulation) that aim to modify it for clinical purposes (Tremblay et al., 2019; Zrenner et al., 2020).

Therefore, understanding the role of oscillations and fast neuronal responses in the interplay between mood and environmental incentives is key and magnetoencephalography (MEG) offers a great opportunity to do so non-invasively.

Here we use a pre-registered approach to identify neural correlates of mood in neuronal oscillations and stimulus evoked responses. This is to do justice to major recent concerns about false positive results in neuroscience in general, but also specifically for results derived from methods with a potentially vast range of features and therefore statistical testing space, as is common in electrophysiology.

In the exploratory analysis of (n= 14, age 16.22 years, described in supplementary materials), we collected MEG data with a monetary gambling task developed by (Keren et al., 2020) to parametrically shift mood by manipulating reward prediction errors (RPEs) and tested with linear mixed effect models whether brain oscillations and trial-level changes in evoked responses would show a relationship with mood or its model components.

This approach allowed us to build and pre-register the following hypotheses which we test in the paper (pre-registered analysis available on OSF (https://osf.io/djw8h), more detailed background on the hypotheses formation is available in the supplementary material):

1. We hypothesize that beta-gamma oscillatory power (25-40Hz) measured by MEG in the time interval preceding mood rating will be positively correlated with reward expectation term derived from the primacy mood model, with a source space cluster covering the frontal superior and medial cortex and the ACC.
2. RPE and self-reported mood will be correlated with the variability in the evoked response to the gamble outcome (feedback), in right precuneus and paracentral lobule (at ~500ms) for RPE and in the right insular cortex (at ~400ms after feedback presentation) for mood.
3. The reward expectation term from the primacy mood model will be predictive of the signal in posterior MEG sensors during the period 250-400ms after presentation of gambling options.

While previous studies have looked at influence of mood on trial-averaged responses (Paul and Pourtois, 2017), to our knowledge this is the first study to relate the temporal dynamics of mood to trial level variations in evoked responses and oscillatory power with non-invasive electrophysiology in humans. Characterizing the neural substrates that link mood and reward is essential to understanding how disrupted reward processing contributes to mood disorders and may be instrumental in predicting symptom trajectory and response to treatment.

## Materials and Methods

### Sample

Subjects are adolescent volunteers (age 12 to 19 years) recruited through mail, online advertisement and direct referrals from clinical sources. Subjects provided informed consent to a protocol approved by the NIH Institutional Review Board before completing questionnaires and an in-person evaluation with a medical practitioner at the NIH clinical center to guarantee their suitability to enroll in the study. Both healthy volunteers (not satisfying criteria for any diagnosis according to DSM-5) and patients with a primary diagnosis of major depression (MDD) or sub-threshold depression were included. All participants received the same scripted instructions for their participation in this study. The full list of inclusion and exclusion criteria is outlined in the supplementary document.

Following in-person screening, MEG data were collected with a 275-channel CTF scanner (272 working channels, sampling at 1200Hz, 3rd order synthetic gradiometer configuration) and a structural MRI (MPRAGE, 1mm isotropic resolution) of the subject’s head was acquired with a 3T GE MRI scanner (collected within 6 months of the MEG scan), both housed in the NMR suite of the NIH clinical center.

We collected MEG data from 56 volunteers that passed our inclusion criteria. Of this sample 2 subjects were excluded from all reported analysis, one due to artifacts during data collection, and one from reporting to have misunderstood task instructions when debriefed at the end of the scanning session. Of the included 54 subjects (age 16.3 ± 1.8 years, 30 MDDs, 35 females), all were included in sensor based analyses, and 51 (age 16.3 ± 1.8 years, 29 MDDs, 33 females) were included in our source space analysis (two subjects did not have a structural MRI due to laboratory shutdown in March 2020 and one subject had large errors >>5mm in the initial localization of the MEG fiducial coils). 14 subjects were initially analyzed as an exploratory sample to inform the study hypotheses (as reported in our preregistration available on OSF (https://osf.io/djw8h)), leaving a separate confirmatory sample of 40 subjects for analyses at the sensor level and a subsample of 37 subjects at the source level.

### Task description

The task consists of three blocks, where in each block a closed-loop mood controller delivers reward prediction errors to try to move the participant’s mood to a target value (see (Keren et al., 2020) for details on the task design). The targets of the controller are to reach the highest (first block), lowest (second block), and then highest mood again (third block). Participants report their mood on a sliding scale with the words “Unhappy” on the left end and “Happy” on the right end of the scale. Within each block 70% of trials are congruent (delivering positive RPE in the high mood target blocks and negative RPE in the low mood block) and 30% are incongruent (delivering negative RPEs in the high mood target blocks and positive RPEs in the low mood block).

Before entering the scanner, participants are instructed on how to perform the task. Participants are told that the final amount they win in the task will be converted to a proportional amount of money, but they are not aware of the mood manipulation.

Each block consists of 27 trials, for a total of 81 trials. An example of a task trial selecting the gambling option is shown in Figure 1. In each trial participants are presented with a certain amount (displayed on the left side of the screen) and two possible gambling outcomes (on the right side of the screen). The gambling amounts are selected by the closed loop controller based on the mood target and the participant’s self-reported mood. Participants are given 3s to decide to either gamble or select the fixed outcome, by pressing the right or left button on a fiber optic response pad (FORP). When participants do not make a selection in time, the task controller selects the gambling option.

**Figure 1:**
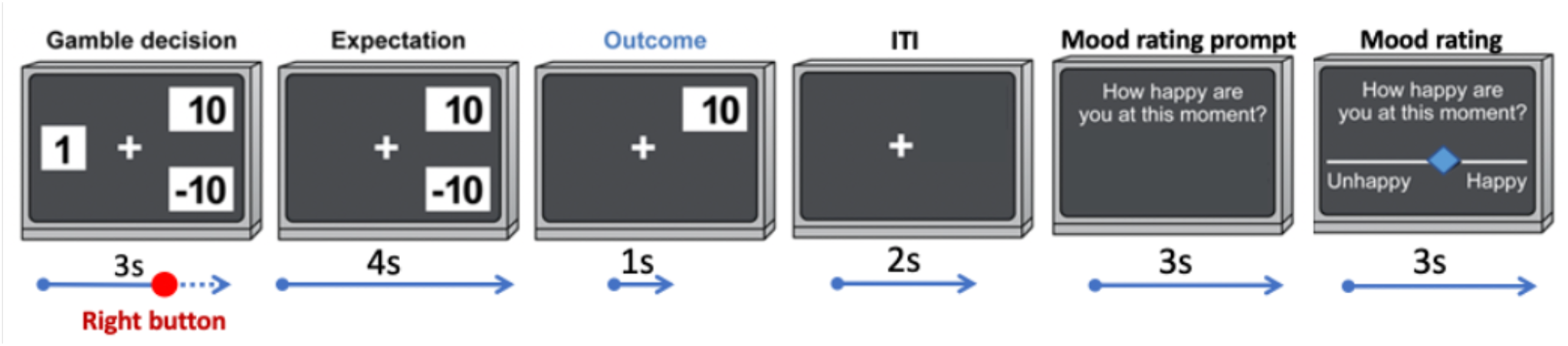
Structure of the closed loop gambling task

After choice selection there is a 4s waiting period, then (for the gamble selection) the gamble outcome is revealed and remains on the screen for 1s. After each trial there is a 2s inter-trial-interval time when only a fixation cross is displayed. Every 2 or 3 trials participants are prompted to rate their current mood on a horizontal slider. The cursor on the slider can be moved continuously by pressing and holding the right and left buttons on the FORP.

At the end of each block participants are given a break from the task while remaining in the scanner and can proceed to the next block by pressing a button on the FORP.

At the end of the MEG scanning session participants are debriefed to ask about their experience in the scanner.

### Mood model

Participants rate their mood with a slider between a value of 0 and 100 (with 0 being the lowest and 100 being the highest) every 2 to 3 trials of the gambling task. Reward prediction error and expectation are estimated according to the primacy mood model proposed by (Keren et al., 2020) defining the mood at time *M*_*t*_ as:

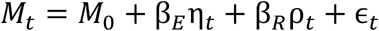

where *M*_0_ is the participant’s baseline mood, β_*E*_ and β_*R*_ represent the sensitivity to expectation and surprise (prediction error) respectively, and ϵ_*t*_ is the error term.

Given a trial outcome value *A*_*t*_ the expectation term from the model is calculated as the average of all previously received outcomes:

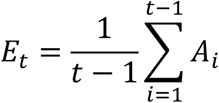

The reward prediction error is then defined as the difference between the outcome and the expectation:

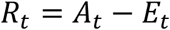

Finally the contributions of expectation and RPE to mood at time *t* are defined as:

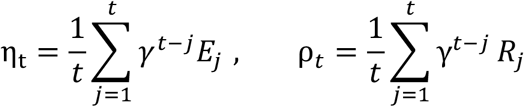

The model parameters M_0_, γ, β_*E*_ and β_*R*_ are derived by fitting the self-reported mood to the mood model with python’s TensorFlow package as described in (Keren et al., 2020).

All trial dependent parameters obtained from the model fit are sampled at every trial, but subjects report mood every 2-3 trials during the gambling task. In order to use self-reported mood in our linear mixed model to test variability over all trials, mood ratings are interpolated with a Piecewise Cubic Hermite Interpolating Polynomial implemented in Matlab.

### MEG data

MEG data analysis is performed on the NIH HPC Biowulf cluster (http://hpc.nih.gov) with Matlab (The MathWorks, Inc., Natick, Massachusetts, United States) and functions from the FieldTrip toolbox ((Oostenveld et al., 2011); http://fieldtriptoolbox.org). MEG data are initially visually inspected and pre-processed with the CTF software: third-order synthetic gradiometer configuration is applied; segments with motion exceeding a threshold of 5mm or including noticeable artefacts are eliminated; data are bandpass filtered between 0.5 and 300Hz, baseline corrected, and a 60Hz notch filter is applied to reduce power line noise. Data are then corrected for eye movements and heartbeat artifacts with ICA fastica algorithm (30 independent components are calculated and a maximum of 4 components, 2 heartbeat and 2 eye movement ICs, are eliminated for each dataset).

### Data Processing

Source data reconstruction is achieved by a beamformer approach and a forward model based on Nolte’s spherical approximation (Nolte, 2003; Veen et al., 1997) implemented in FieldTrip with subjects’ individual brain MRIs co-registered to the MEG data. Source level activity is reconstructed on a 5mm grid based on the MNI brain and warped to individual anatomy. Beamformer weights are calculated based on the data covariance over the whole task with a covariance regularization equal to 5% of its maximum singular value (a high regularization is selected to improve SNR and increase spatial smoothness (Brookes et al., 2008); the effects of matrix regularization and forward model selection is explored in supplementary analysis).

### Beta-gamma power

Following our pre-registered hypotheses, we estimate beta-gamma oscillatory power in the 3s waiting period preceding mood-rating. MEG signal is first band pass filtered in the frequency band of interest (25-40Hz) then, for sensor space analysis oscillatory power is estimated by measuring the signal variance in each 3s window. For source space analysis oscillatory power in each voxel *i* is estimated as

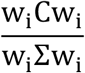

where w_i_ are the beamformer weights, C is the data covariance matrix in the 3s window of interest, and Σ the estimated noise covariance matrix.

### Evoked responses

For the analysis of evoked responses, a low-pass filter of 30Hz is applied following the pre-processing steps before calculation of the beamformer weights.

- Gamble feedback Regions of interest (ROI) are defined by the Automated Anatomical Labelling (AAL) atlas available in fieldtrip. The signal from an ROI is estimated from its geometrical centroid. We select a time window −200 to 1000 ms with respect to the presentation of the gamble feedback. The source data time course is estimated and then downsampled to 300Hz to reduce the number of time points to test.
- Gamble option We select the average response in the time window of 250 to 400 ms with respect to the presentation of the gambling options for the analysis of the evoked responses at both sensor and source level. For source space analysis the data time course is first reconstructed at each source voxel and then the evoked response for each trial is estimated as the average signal in the 250 to 400ms time window. Regions of interest (ROI) are defined by the Automated Anatomical Labelling (AAL) atlas available in fieldtrip. The signal from an ROI is estimated from its geometrical centroid. We select a time window −200 to 1000 ms with respect to the presentation of the gamble feedback. The source data time course is estimated and then downsampled to 300Hz to reduce the number of time points to test.

It is important to note that with beamforming the sign of evoked responses in source space is uncertain (i.e. sign may be flipped between participants). For each participant the sign of the source signal for an ROI or voxel is estimated by maximizing the correlation over subjects of their trial averaged evoked response to the stimulus (gambling option or feedback presentation).

### Statistical Analysis

We apply linear mixed effects models to estimate the contribution of trial level mood, expectation and reward prediction error to response variability in the MEG data. The use of linear mixed effect models allows us to analyze data from all participants in a single model while accounting for inter-subject differences.

### Hypothesis 1

The effect of expectation on beta-gamma power is tested for 2 separate predictors *E*_*t*_ and η_*t*_.

Our formulation is as follows (here presented for the *mood* fixed effect):

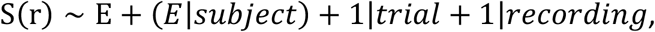

where S(r) indicates the MEG signal at location r (either sensor or source voxel). S(r) is a vector with dimension 1 × N_m_, where N_m_ is the number of mood rating trials from all subjects.

### Post-hoc analyses

The following post-hoc analyses are run with the whole sample of available subjects (N=51 at the source level).

### Mediation analysis

We hypothesize that beta-gamma power variability over trials is related to reward expectation, as defined by the primacy mood prediction model. Expectation, by its definition (see equation 1) is a predictor of self-reported mood. In order to test if self-reported mood is at all directly related to beta-gamma power we run a mediation analysis with beta-gamma power as the independent variable, the expectation term as the mediator variable and mood as the dependent variable.

### Comparison with previous fMRI results

Keren et al. previously published results using the same gambling task and primacy mood model with fMRI data. They reported a significant cluster of BOLD activation in the ACC correlating with the subject specific expectation weight, β_*E*_ (see equation 1). Following our hypothesis that trial variations in beta-gamma power are related to reward expectation we then test if the subject average beta-gamma power is correlated to β_*E*_ in an analogous analysis to (Keren et al., 2020). For this analysis the average beta-gamma power is calculated at the voxel level by averaging the previously estimated MEG power in the 3s pre mood rating period over all available trials for each subject.

A Pearson correlation between subject average beta-gamma power and β_*E*_ is then run and significant clusters are calculated. Null distributions are obtained by calculating maximum TFCE value in random permutations (N=5000).

The significant cluster from Keren et al. is then compared to the MEG cluster in MNI space to check for congruent cross-modality results.

### Hypothesis 2

In order to test the effect of mood and RPE on the evoked response following gambling feedback we run models independently for 3 separate fixed effects: self-reported mood, trial RPE as defined in the primacy model (R_)_) and the primacy weighted RPE term (ρ_)_). Our formulation is as follows (here presented for the *mood* fixed effect):

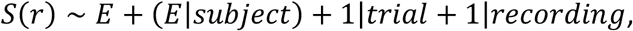

where S(t, r) indicates the source reconstructed MEG signal in ROI *r* and at time *t* with respect to the task event. *S*(*t, r*) is a vector with dimension 1 × *N*_*f*_, where *N*_*f*_ is the number of feedback trials from all subjects.

In our confirmatory analysis we test if mood predicts MEG signal in the right insula, and if the two RPE parameters predict response variation in the right paracentral lobule and precuneus as defined by the AAL atlas.

In addition, as a post-hoc analysis, the data from all available subjects is combined (exploratory + confirmatory sample) and each predictor is tested on all 116 ROIs from the AAL atlas. For this post-hoc analysis, temporal clusters are corrected for multiple comparisons over all ROIs and the 3 predictors.

### Hypothesis 3

We test the effect of trial expectation, as defined by the primacy model (*E*_*t*_), and the primacy weighted expectation term (η_*t*_) on the average evoked response in the 250-400ms window following gamble options presentation.

Our formulation is as follows (here presented for the *E* fixed effect):

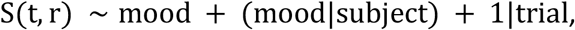

where *S*(*r*) indicates the MEG signal at location *r* (either sensor or source voxel).

*S*(*r*) is a vector with dimension 1 × *N*_o_, where *N*_o_ is the number of task trials from all subjects.

We run our statistical model at each sensor and source space voxel.

For all linear mixed effect models we test for statistical significance over multiple voxels or sensors by applying Threshold Free Cluster Enhancement (TFCE) spatial clustering with parameters E=0.5, H=2, dh = 0.1 as indicated in (Smith and Nichols, 2009).

Null distributions are obtained by running the same linear mixed model after random permutation of S over trials for each subject (10,000 random permutations will be used for the sensor space analysis, 2,000 random permutation will be used for the voxel space). Each random permutation is used for all sensors/voxel, TFCE is applied on the t-values for the fixed effect and the maximum (and minimum) spatial cluster value is included in the null distribution.

We then use a 2-tailed t-test against the null distribution to infer statistical significance of the spatial clusters (α = 0.05) corrected with false discovery rate for the multiple fixed effects we are testing.

## Results

### Hypothesis 1: trial variations in beta-gamma oscillatory power measured in the 3s time interval preceding mood rating will be positively correlated with *E*_*t*_

For our first hypothesis we set to test if trial variations in beta-gamma (25-40Hz) oscillatory power are affected by reward expectation.

We report results for the linear mixed model analysis with trial expectation (*E*_*t*_) and primacy weighted expectation (η_*t*_) as fixed effects of the model. At the sensor level we found a significant cluster of sensors (Figure 2), for both *E*_*t*_ (peak on channel MRC22, fixed effect t-stat = 3.94, uncorrected p-value=8.6e-05) and η_*t*_ (peak on channel MRC25, fixed effect t-stat = 3.64, uncorrected p-value=2.8e-04).

**Figure 2:**
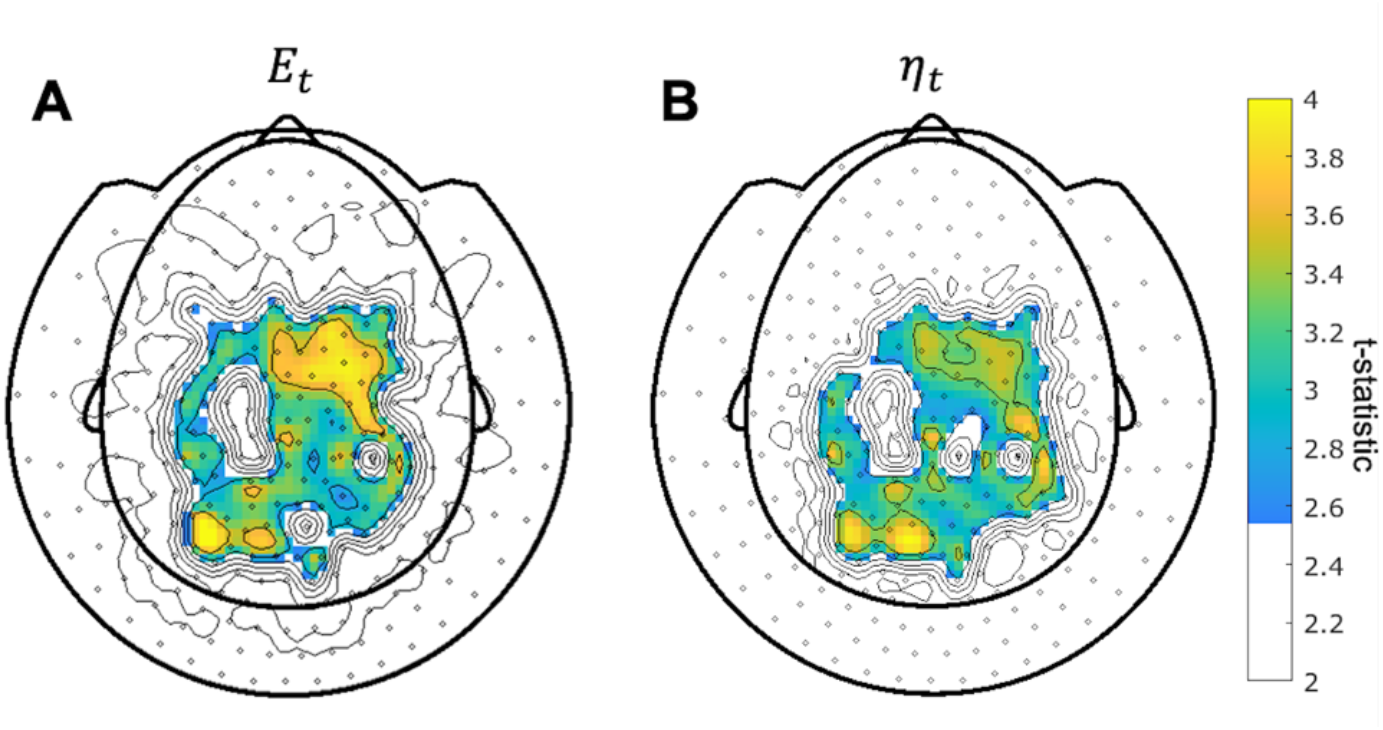
Maps of MEG sensors where expectation parameters E_t_ (A) and η_t_ (B) significantly predict beta-gamma power preceding mood rating in confirmatory sample. Sensors surviving clustering correction (α < 0.05, two-tailed, 10,000 random permutations) are shown in color. Color bar indicates t-statistc of the fixed effect.

At the source level we found a significant cluster where *E*_*t*_ predicted beta-gamma power in the mid to posterior cingulate cortex and extending to the paracentral lobule (Figure 3A, cluster peak at MNI coordinate [−2, −40, 34]mm, T-stat=5.21, uncorrected p-value=2.3e-07). A similar, but less significant cluster appeared for η_*t*_ (Figure 3B cluster peak at MNI coordinate [−2, −40, 30]mm, T-stat=4.78, uncorrected p-value=2.0e-06). Other significant clusters were present in the occipital cortex, caudate and the ACC. By exploration of the covariance regularization parameter (5%, 1% and 0.2% of the maximum singular value of the covariance matrix) we found significant clusters to be highly dependent on regularization, with clusters in supplementary motor cortex and frontal superior cortex becoming prominent at lower regularization values (Figure S4 in supplementary material).

**Figure 3:**
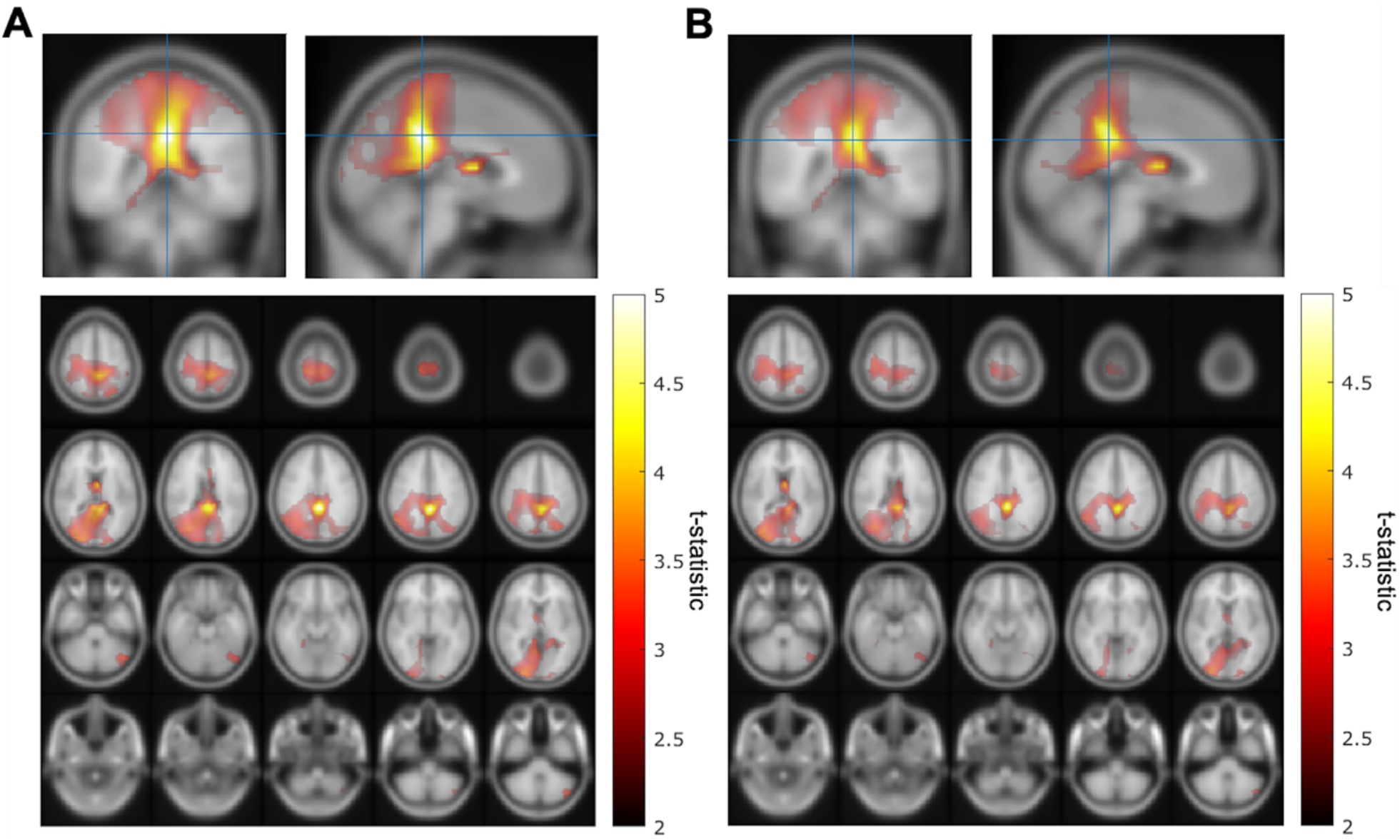
Source space maps showing brain regions where expectation parameters E_t_ (A) and η_t_ (B) predict beta-gamma power preceding mood rating in confirmatory sample. Color bar indicates t-statistic of the fixed effect. Top row shows cluster peak in coronal and sagittal views. The plot on the bottom shows the significant cluster over multiple axial slices. Both predictors show similar significant clusters with a peak in the left posterior cingulate cortex (MNI coordinate [−2, −40, 34]mm for E_t_ and [−2, −40, 30]mm for η_t_) extending to mid cingulate, parietal cortex and the caudate.

Over our explored range of regularization values the mid-posterior cingulate cortex cluster remained present.

We found that subject average beta-gamma power in the pre mood rating period was significantly correlated with the subject expectation weight from the primacy mood model (Figure 4C,D). We found significant clusters with peaks in the ACC, caudate and occipital cortex. By comparing Keren et al. (Keren et al., 2020) previous fMRI work we found overlap in the ACC cluster between MEG average beta-gamma power and BOLD fMRI activation.

**Figure 4.**
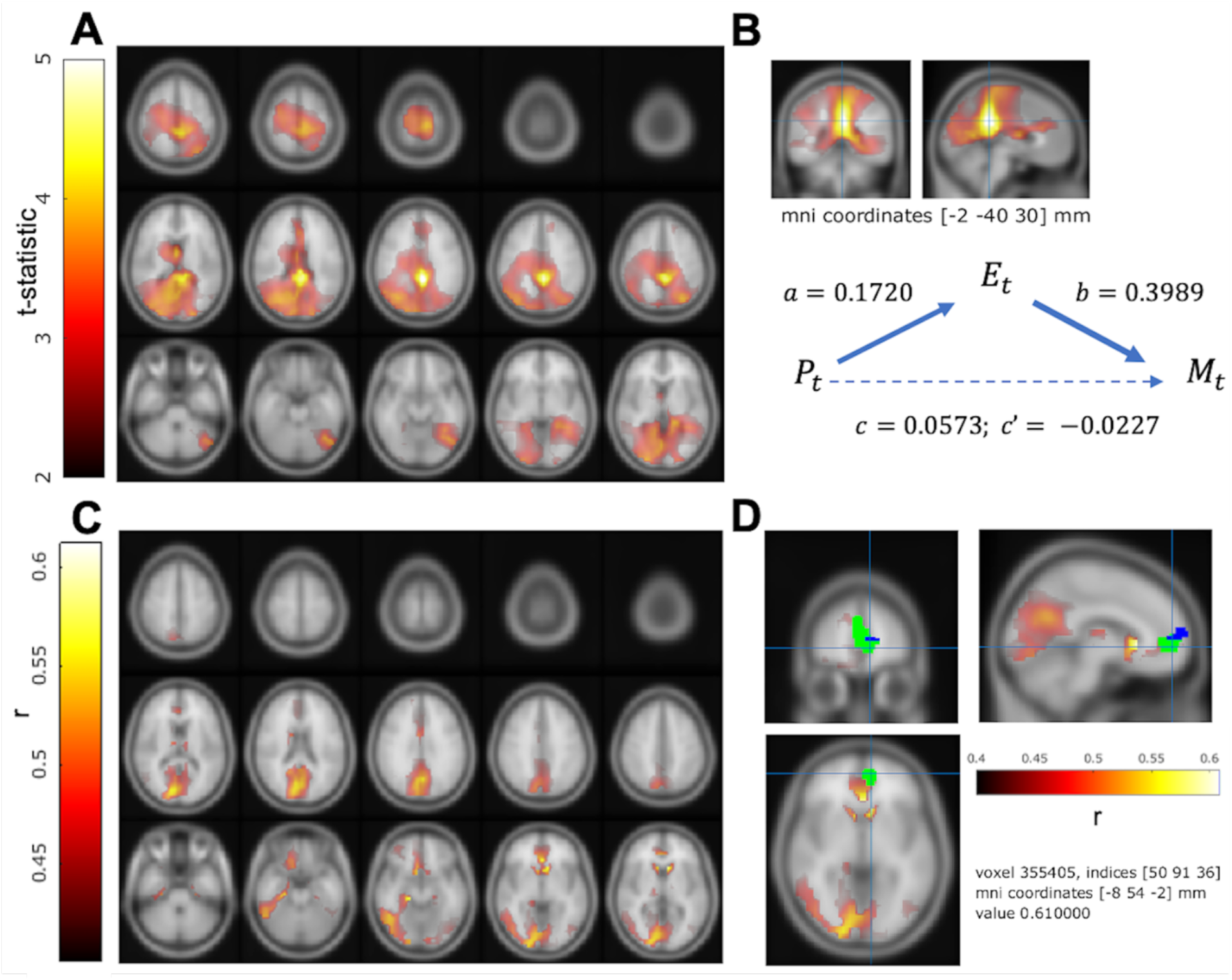
Post-hoc analysis of beta-gamma power with all available subjects (N=51). **A**. Trial-level analysis: source space maps showing brain regions where expectation parameters E_t_ predicts beta-gamma power preceding mood rating (see confirmatory results in figure 3A). Color bar indicates t-statistic of the fixed effect. **B**. Mediation model where the reward expectation E_t_ mediates the relationship between beta-gamma power P_t_ and subject’s self-reported mood M_t_. At the source cluster peak we found evidence of complete (ab/c = 1.1973) inconsistent mediation (significant indirect effect ab = 0.0686 ± 0.018 (Z = 3.6271)). **C**. Subject level analysis: axial view of regions where the subject level expectation weight β_E_ significantly (α < 0.05, 5,000 random permutations) correlates with subject average beta-gamma power preceding mood rating. Color bar indicates Pearson’s correlation (r), source map has been masked with cortical and subcortical regions included in the AAL atlas. Subject beta-gamma power shows significant correlation clusters in subgenual ACC, caudate and occipital cortex. **D**. Regions where β_E_ correlates with brain activity in MEG (**C**) and fMRI (significant region from Keren et al.). MEG source localization is displayed in the heat color map, voxels of overlap between MEG and fMRI results are displayed in green and voxels where only fMRI showed significant correlation are in blue.

In order to test if beta-gamma power is reflective of subject’s mood we tested whether this relationship could be mediated by our expectation parameter. We found that indeed reward expectation *E*_*t*_ significantly mediated (significant indirect effect ab = 0.0686 ± 0.018 (Z = 3.6271)) the relationship between beta-gamma power *P*_*t*_ and subject’s self-reported mood *M*_*t*_ with a complete (ab/c = 1.1973) inconsistent mediation (Figure 4B).

### Hypothesis 2: self-reported mood and RPE from the primacy model can predict changes in the response to reward. *R*_*t*_ and *ρ*_*t*_ will be correlated with the variability in the evoked response in right precuneus and paracentral lobule (at ~500ms) and for *mood*_*t*_ in the right insular cortex (at ~400ms after feedback presentation)

In order to test the effect of mood and RPE on the evoked response following gambling feedback we ran models independently for the three separate fixed effects. From our exploratory analysis we hypothesized to find significant clusters in the right insula cortex for the mood predictor and right precuneus and paracentral lobule for *R*_*t*_ and ρ_*t*_. In our confirmatory analysis we found no significant effect of self-reported mood on the evoked response source localized to the right insula.

Both *R*_*t*_ and primacy weighted ρ_*t*_ parameter were confirmed to predict the response on the right paracentral lobule at 500ms, with ρ_*t*_ also predicting an earlier peak at ~250ms (Figure 5, Table 1). Neither parameter predicted the response in the right precuneus at 500ms, but we saw a significant effect of *R*_*t*_ on an earlier peak (100ms).

**Table 1:** Summary of results from the confirmatory analysis for hypothesis 2 (time courses in figure 5). We tested the responses to reward feedback presentation in the right insula to self-reported mood, and the right precuneus and right paracentral lobule to reward expectation.

| ROI | Predictor | Window(ms) | Peak time(ms) | T-stat | p-value |
| --- | --- | --- | --- | --- | --- |
| Insula Right | mood | Not significant |  |  |  |
| Precuneus Right | R | 97-111 | 101 | -3.98 | 7.1e-05 |
| | $\rho$ | Not significant | | | |
| Paracentral Lobule Right | R | 472-502 | 489 | 3.33 | 8.7e-04 |
| | $\rho$ | 225-278 | 248 | -4.33 | 1.5e-05 |
| | $\rho$ | 465-515,<br>629-639 | 502 | 3.13 | 1.8e-03 |

**Figure 5:**
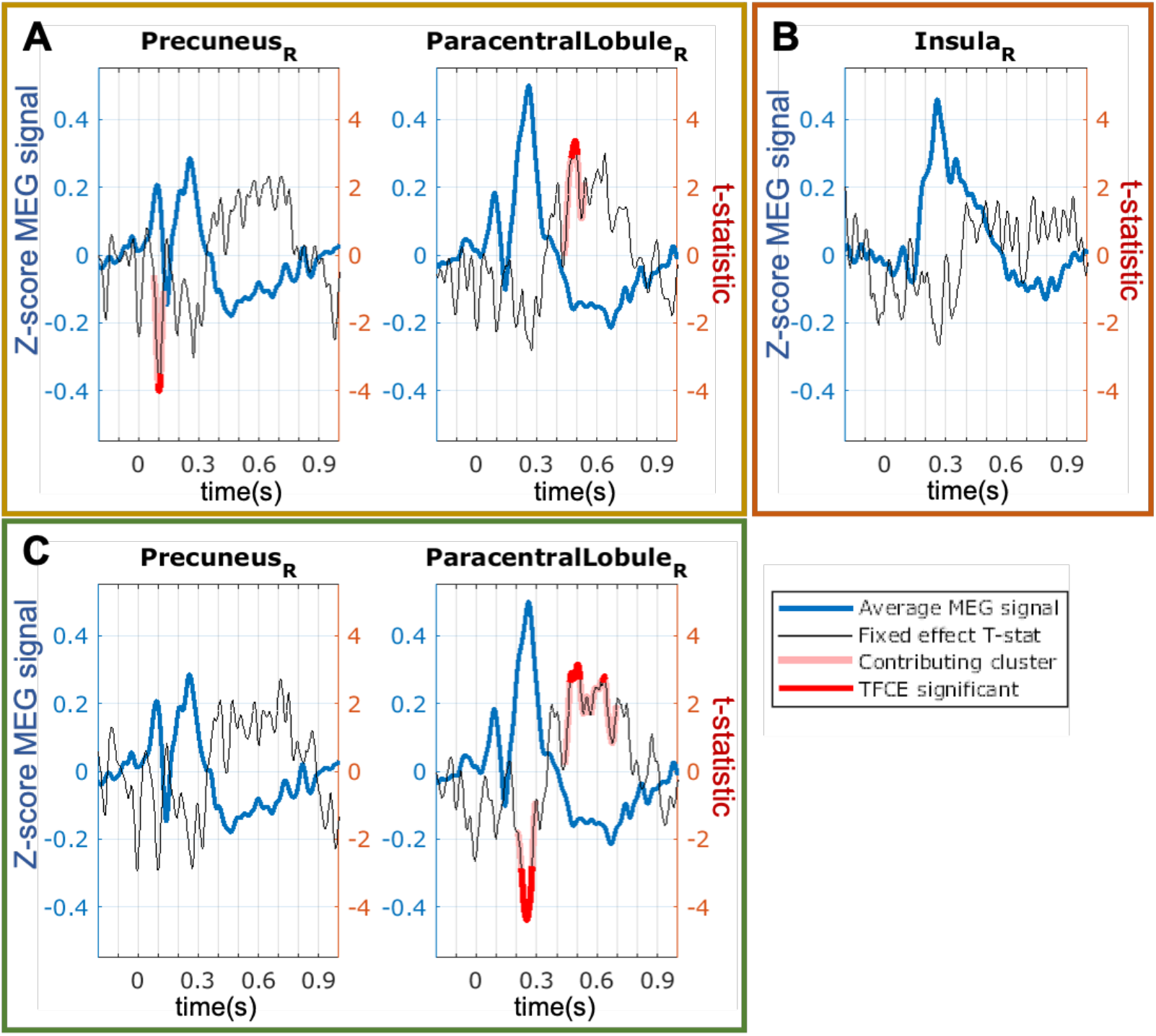
Response to reward feedback in regions of interest (ROIs) from the AAL atlas hypothesized to vary with mood or with the reward prediction error parameters. **A**. Response to feedback in the right insula cortex and prediction from self-reported mood. No significant temporal clusters were found in the confirmatory sample. **B**. ROIs with significant temporal clusters to the R_t_ fixed effect. Average response is in blue, t-statistic of the fixed effect is in black and significant temporal clusters are in red. R_t_ significantly predicts trial level variations in reward feedback evoked response in right precuneus (cluster peak at 101ms, not supporting our initial hypothesis) and right paracentral lobule (cluster peak at 489ms, in agreement with hypothesis). **C**. ROIs with significant temporal clusters to the ρ_t_ fixed effect. ρ_t_ was confirmed to predict responses in right paracentral lobule (cluster peaks at 248ms and 502ms). No significant effect of ρ_t_ was found in the right precuneus.

From our further post-hoc analysis we found that both *R*_*t*_ and ρ_*t*_ significantly predicted the response in the left angular cortex (peak at ~250ms). Moreover the ρ_*t*_ parameter significantly predicted a late response (~600ms) in the left paracentral lobule and a peak in the right para hippocampal gyrus (~430ms) (Figure 6, Table 2). Self-reported mood did not significantly predict the response to reward feedback in any ROI.

**Table 2:**
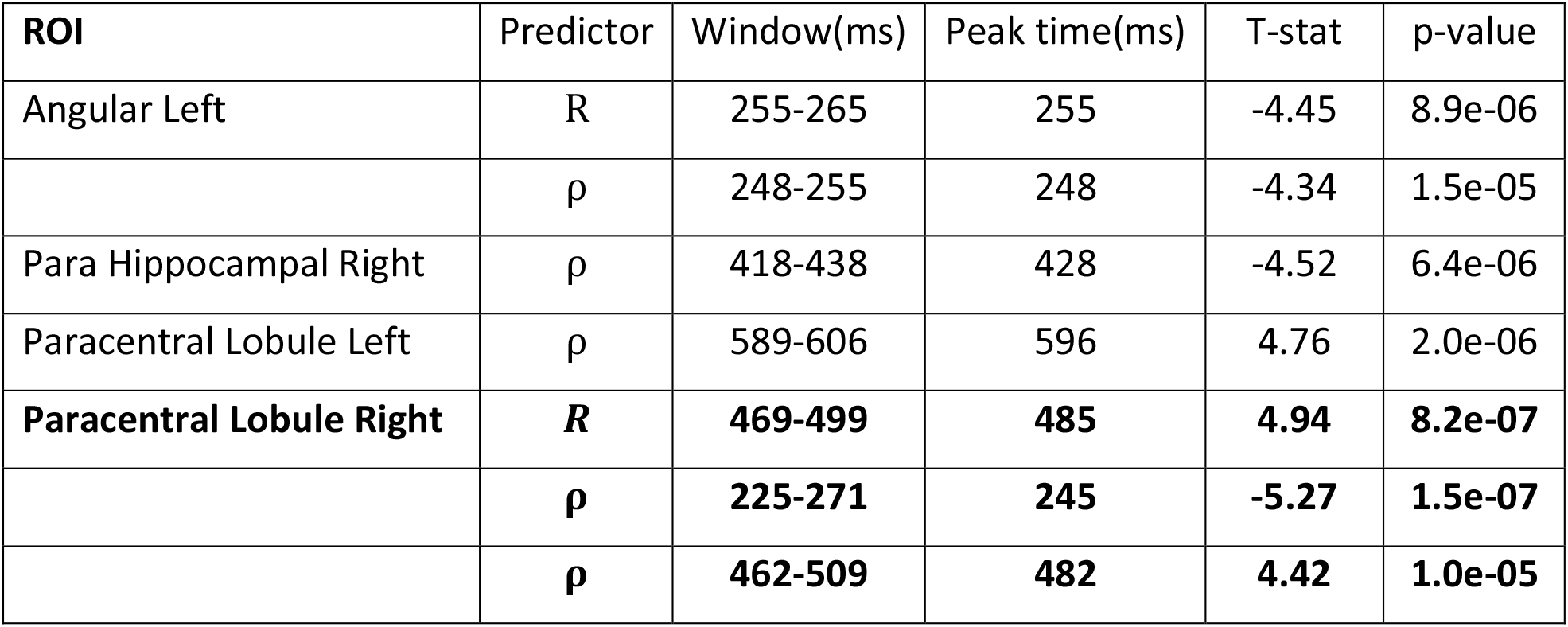
Summary of results from the post-hoc analysis over all ROIs on responses to reward feedback presentation (time courses in figure 6). Significant peaks in the right paracentral lobule, highlighted in bold, are also seen in the confirmatory analysis (table 1).

| ROI | Predictor | Window(ms) | Peak time(ms) | T-stat | p-value |
| --- | --- | --- | --- | --- | --- |
| Angular Left | R | 255-265 | 255 | -4.45 | 8.9e-06 |
| | $\rho$ | 248-255 | 248 | -4.34 | 1.5e-05 |
| Para Hippocampal Right | $\rho$ | 418-438 | 428 | -4.52 | 6.4e-06 |
| Paracentral Lobule Left | $\rho$ | 589-606 | 596 | 4.76 | 2.0e-06 |
| **Paracentral Lobule Right** | **R** | **469-499** | **485** | **4.94** | **8.2e-07** |
|  | **$\rho$** | **225-271** | **245** | **-5.27** | **1.5e-07** |
|  | **$\rho$** | **462-509** | **482** | **4.42** | **1.0e-05** |

**Figure 6.**
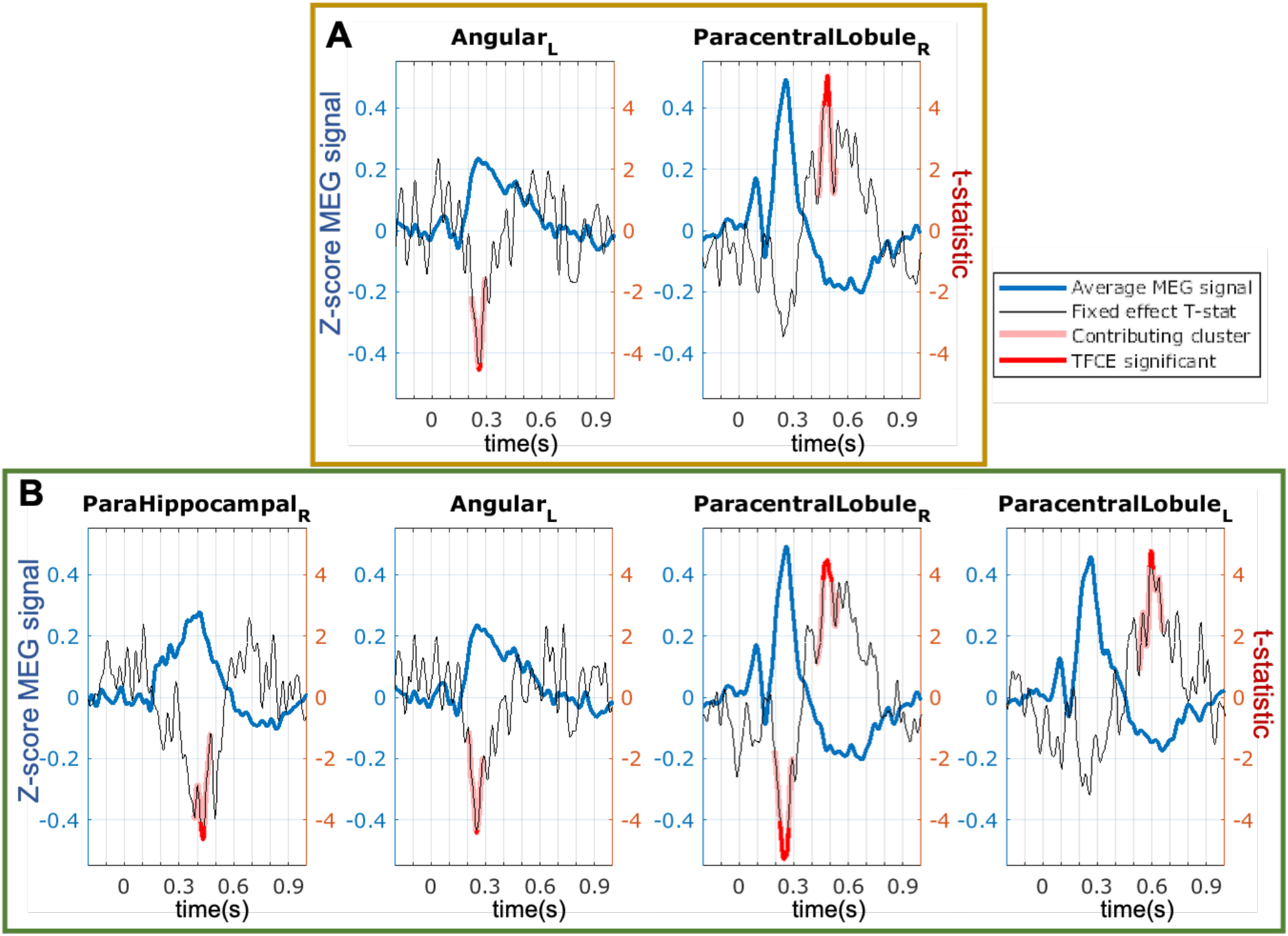
Regions of interest (ROIs) from the AAL atlas significantly predicted by reward prediction error in post-hoc analysis (including all available subjects, N=51). **A**. ROIs with significant temporal clusters to the R_t_ fixed effect. Average response is in blue, t-statistic of the fixed effect is in black and significant temporal clusters are in red. **B**. ROIs with significant temporal clusters to the ρ_t_ fixed effect. In addition to the confirmed effect on the right paracentral lobule (see figure 5), we found that both parameters predicted trial variations in the left angular gyrus and ρ_t_ in the right para-hippocampal gyrus and left paracentral lobule.

### Hypothesis 3: the reward expectation parameters from the primacy mood model (*E*_*t*_ and *η*_*t*_) will be predictive of the signal in posterior MEG sensors during the 250-400ms time window after the presentation of gambling options

For our last hypothesis we tested whether the trial level variation in evoked response 250-400ms after presentation of gambling options could be predicted by the model expectation parameters. Agreeing with our initial hypothesis we found that both *E*_*t*_ and η_*t*_ significantly predicted signal response in clusters of MEG axial gradiometers (figure 7D). At the source level we found a significant cluster in the right cerebellum (figure 7E) only for the *E*_*t*_ predictor.

**Figure 7:**
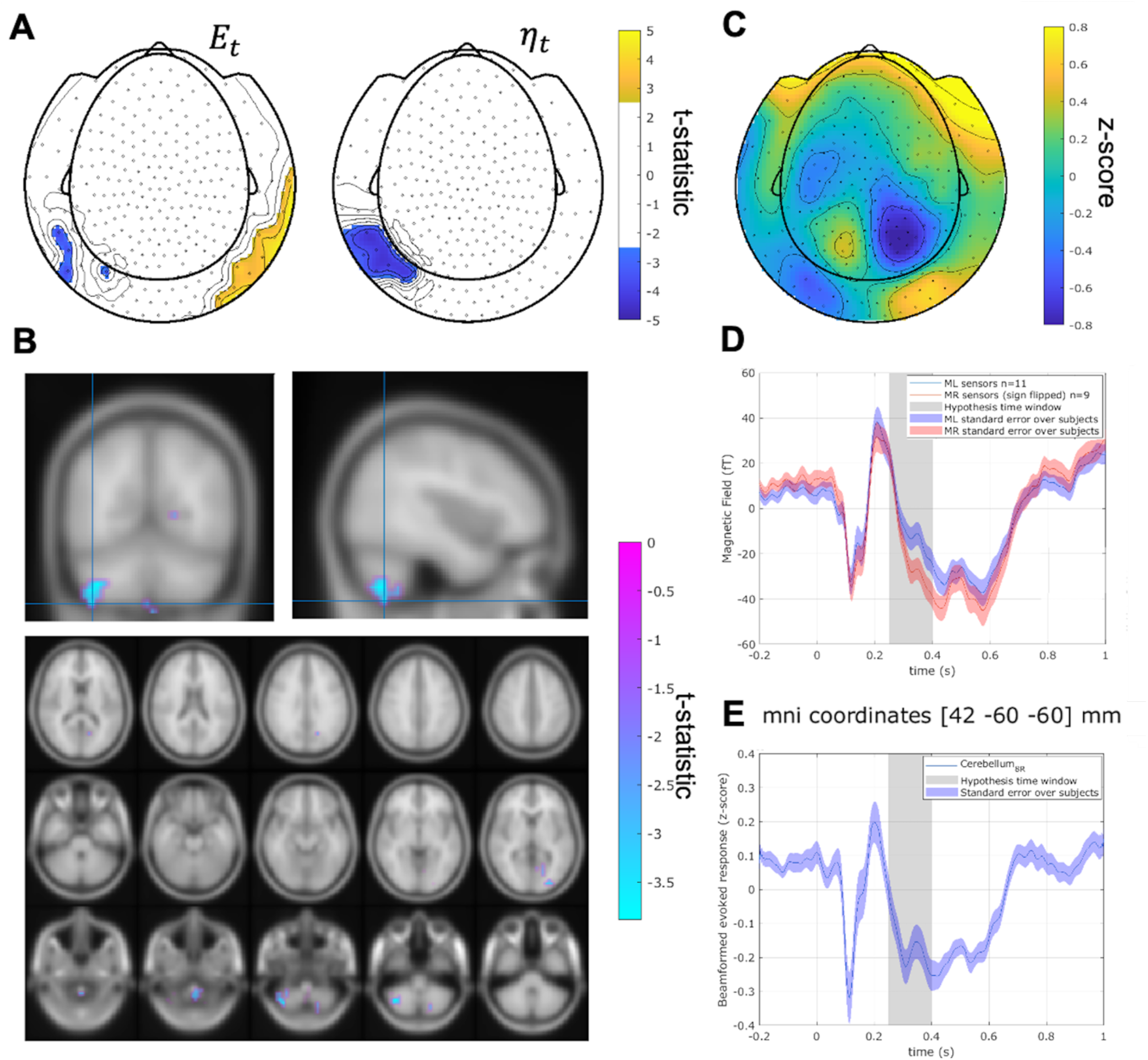
Expectation predicts the evoked response 250-400ms after presentation of gambling options. **A**. Maps of MEG sensors where expectation parameters E_t_ (left) and ρ_t_ (right) significantly predict MEG response 250-400ms following presentation of gambling options. Sensors surviving clustering correction (α < 0.05, two-tailed, 10,000 random permutations) are shown in color. Color bar indicates t-statistic of the fixed effect. **B**. Source space clusters where expectation parameters E_t_ predict MEG signal response. Predictor ρ_t_ did not show any significant clusters. Color bar indicates t-statistic of the fixed effect. Top row shows cluster peaks in coronal and sagittal views. The plots on the bottom show the significant clusters over multiple axial slices. The expectation parameter was predictive of activity in the right cerebellum. **C**. Topographic map of the z-scored average MEG signal over all trials and subjects in the 250-400ms time window after presentation of gambling options. **D**. Time course of the evoked response (average over all trials and subjects in confirmatory sample) over all significant sensors (clusters in figure 7A). The average of the left sensors (n=11) is in blue. The sign of the right sensors (n=9) in red has been flipped for easier comparisons. The time window of interest is highlighted in grey. **E**. Time course of the evoked response (average over all trials and subjects in confirmatory sample) in the significant area of the right cerebellum from figure 7B.

## Discussion

With a confirmatory sample of 40 participants, we found support in our pre-registered hypothesis that reward expectation, defined by the computational primacy mood model, is positively related to beta-gamma power over central MEG sensors. The same linear mixed effect analysis at the source space analysis localized the main significant cluster over the posterior cingulate cortex (PCC), extending to mid cingulate and paracentral lobule and parietal cortex. Unlike the exploratory sample no clusters were found in superior or medial frontal cortex. The cingulate cortex is thought to have an important role in integrative brain functions, being involved in emotional processing, memory and learning. The PCC in particular is involved in memory processes and strongly connected to the hippocampus as well as being a key node in the resting state network.

It is interesting to note that from our model definition, reward expectation is the equivalent to the average of all experienced rewards during the gambling task, a mental representation that we expect to involve memory processes. Our results do not link directly beta-gamma power with self-reported mood, but to reward expectation, which previous work has found to be a key component in mood processes. Through a post-hoc mediation analysis we found that reward expectation fully mediates any relationship between mood and beta-gamma power, meaning that self-reported mood by itself would not explain any variation in the same MEG data. This highlights how it may be important to identify the separate mental processes that conflate to shift mood over time if we hope to understand the integrative function of mood as a whole.

With this result we also identify that brain oscillations may have an important, and measurable, role in mood dynamics, as previously found in studies involving pharmacological manipulation (e.g. ketamine) and group comparisons between MDDs and HVs showing in particular how reduced frontal gamma oscillations may be a marker of depression (Fitzgerald and Watson, 2018; Nugent et al., 2019a).

Brain oscillations are believed to be key to brain communication (supported by an expanding literature in MEG functional connectivity) and the function of brain oscillations is dependent on frequency, which in conjunction with other techniques (PET, MRS), can be used to explore the specific function of neuronal assemblies (e.g. glutamatergic excitatory vs GABAergic inhibitory processes). This is key to determine possible dysfunctions in mood disorders such as depression, and possibly developing more effective pharmacological therapies than current antidepressants. Beta-gamma oscillations have been observed in multiple EEG studies, synchronizing in response to positive feedback (Hosseini and Holroyd, 2015; Marco-Pallarés et al., 2015). We initially selected the 25-40Hz band based on these studies and following test in our exploratory sample with standard frequency bands. The naming of the band in the literature is unclear, with different authors also referring to it as high-beta or low-gamma bands. It is worth exploring if changes in beta-gamma power can be observed in direct response to reward presentation (as observed in EEG following reward feedback) and their precise localization through different MEG task designs or by combination with fMRI techniques (for example with Representation Similarity Analysis (RSA)).

We found confirmatory evidence that RPE (a predictor of mood as defined from the primacy model) modulates the evoked potential in the paracentral lobule. We had hypothesized that the response of the insular cortex to reward feedback would be correlated with self-reported mood.

The insular cortex is thought to be a key area for reward processes, interoception and mood (Preuschoff et al., 2008; Singer et al., 2009) with multiple studied identifying changes in insula function in depressed patients. While we observed activation of the right insular cortex following reward feedback (both positive and negative), in this study we could not find significant evidence that insula response is directly influenced by participants’ self-reported mood.

In a post-hoc analysis testing the response to reward feedback of all ROIs we found evidence that following presentation of gambling feedback, reward prediction error affects even early responses in the left angular gyrus between 200-300ms (part of the visual stream, contralateral to side of the screen where feedback is displayed).

We did not find an evoked response changing with RPE equivalent to the feedback related negativity, extensively observed in EEG. (Doñamayor et al., 2012) localized the FRN from the MEG data to the PCC. Other studies combing EEG and fMRI localized the FRN to the dorsal ACC (Hauser et al., 2014). Evoked responses are affected by both reward magnitude and expectation. It is likely that the relatively low number of negative feedback trials and high variance of reward amounts presented to each subject might have affected this.

We confirmed our pre-registered hypothesis and found evidence that cerebellum activity is correlated with reward expectation 250-400ms following the presentation of gambling options. Several papers have highlighted the involvement of cerebellum in reward processing and its connection to basal ganglia (Bostan et al., 2010; Pierce and Péron, 2020; Wagner et al., 2017).

Although we tried to mitigate statistical bias by pre-registering our approach, our analysis still has some technical limitations. We found evidence that beta-gamma power is positively correlated with expectation at the sensor level and again at the source level. This confirms our initial hypothesis we set with the pre-registration. We want to point out that we are analyzing the change of oscillatory power over trials: this gives lower signal to noise compared to a standard beamformer localization where multiple trials are averaged together. We are compensating from this by using a linear mixed effects model to include all our available data into one statistical test. This still doesn’t obviate to the determination of beamformer weights (which are highly dependent on regularization and choice of forward model as can be seen in our post-hoc analysis in supplementary material).

Uncertainty in the source localization is due to the ill posed nature of the inverse problem in M/EEG: source localization is an estimate dependent on our head model, signal SNR and co-registration accuracy (Jaiswal et al., 2020).

We suggest that for applying a similar analysis technique in future it might be beneficial to apply an adaptive regularization as proposed in (Woolrich et al., 2011) or test localization accuracy at similar SNR level with computational models to determine the best beamformer parameters.

We can see how key features of rewards that influence mood dynamics (i.e. expectation, and reward prediction error) are encoded in multiple brain regions at different times and with different mechanisms (both stimulus evoked responses and oscillations).

To our knowledge this paper offers new evidence that it is possible to track the effect of changes in mood predictors in neuronal activity (non-invasively) at the minute time scales. Based on our pre-registered exploratory analysis, within our task we only expected to see direct correlates of mood on neural activity on the insula, but this was not confirmed by our results. While univariate analysis did show significant effects of mood, multivariate analysis (like mediation) may be more suited to reveal of our perceived mood affects brain function. We believe mood to be an integrative function, which cannot be accurately reflected by the activity of a single brain area but is the result of activity and communication between multiple cortical and sub-cortical regions. Mood may not be measured as activation of a brain region, but rather a shift in baseline activity and functional connectivity of brain networks, priming the brain to respond more strongly to certain stimuli (and/or “inhibiting” the brain to respond less strongly to others), similar to the effect of attention and arousal (Bowrey et al., 2017). We hope this start may help laying some of the groundwork to determine possible causality in reward and mood processes in humans in vivo by identifying both ROIs and accurate timing.

## Acknowledgments

This research was supported by the Intramural Research Program of the National Institute of Mental Health, National Institutes of Health (NIH) (Grant No. ZIA-MH002957-01 to AS). The funder had no role in the design and conduct of the study; collection, management, analysis, and interpretation of the data; preparation, review, and approval of the manuscript; or decision to submit the manuscript for publication. The views expressed in this article do not necessarily represent the views of the National Institutes of Health, the Department of Health and Human Services, or the United States Government.

This work used the computational resources of the NIH HPC (high-performance computing) Biowulf cluster (http://hpc.nih.gov). Data analysis uses functions from the FieldTrip software toolbox (http://fieldtriptoolbox.org).

## References

Bostan, A.C., Dum, R.P., Strick, P.L., 2010. The basal ganglia communicate with the cerebellum. Proc. Natl. Acad. Sci. 107, 8452–8456. https://doi.org/10.1073/pnas.1000496107

Bowrey, H.E., James, M.H., Aston-Jones, G., 2017. New directions for the treatment of depression: Targeting the photic regulation of arousal and mood (PRAM) pathway. Depress. Anxiety 34, 588–595. https://doi.org/10.1002/da.22635

Brookes, M.J., Vrba, J., Robinson, S.E., Stevenson, C.M., Peters, A.M., Barnes, G.R., Hillebrand, A., Morris, P.G., 2008. Optimising experimental design for MEG beamformer imaging. NeuroImage 39, 1788–1802. https://doi.org/10.1016/j.neuroimage.2007.09.050

De Pascalis, V., Strippoli, E., Riccardi, P., Vergari, F., 2004. Personality, event-related potential (ERP) and heart rate (HR) in emotional word processing. Personal. Individ. Differ. 36, 873–891. https://doi.org/10.1016/S0191-8869(03)00159-4

Doñamayor, N., Schoenfeld, M.A., Münte, T.F., 2012. Magneto- and electroencephalographic manifestations of reward anticipation and delivery. NeuroImage 62, 17–29. https://doi.org/10.1016/j.neuroimage.2012.04.038

Fingelkurts, Alexander A., Fingelkurts, Andrew A., 2015. Altered Structure of Dynamic Electroencephalogram Oscillatory Pattern in Major Depression. Biol. Psychiatry, Cortical Oscillations for Cognitive/Circuit Dysfunction in Psychiatric Disorders 77, 1050–1060. https://doi.org/10.1016/j.biopsych.2014.12.011

Fitzgerald, P.J., Watson, B.O., 2018. Gamma oscillations as a biomarker for major depression: an emerging topic. Transl. Psychiatry 8, 1–7. https://doi.org/10.1038/s41398-018-0239-y

Hauser, T.U., Iannaccone, R., Stämpfli, P., Drechsler, R., Brandeis, D., Walitza, S., Brem, S., 2014. The feedback-related negativity (FRN) revisited: New insights into the localization, meaning and network organization. NeuroImage 84, 159–168. https://doi.org/10.1016/j.neuroimage.2013.08.028

Hosseini, A.H., Holroyd, C.B., 2015. Reward feedback stimuli elicit high-beta EEG oscillations in human dorsolateral prefrontal cortex. Sci. Rep. 5. https://doi.org/10.1038/srep13021

Huang, M.X., Mosher, J.C., Leahy, R.M., 1999. A sensor-weighted overlapping-sphere head model and exhaustive head model comparison for MEG. Phys. Med. Biol. 44, 423–440. https://doi.org/10.1088/0031-9155/44/2/010

Jaiswal, A., Nenonen, J., Stenroos, M., Gramfort, A., Dalal, S.S., Westner, B.U., Litvak, V., Mosher, J.C., Schoffelen, J.-M., Witton, C., Oostenveld, R., Parkkonen, L., 2020. Comparison of beamformer implementations for MEG source localization. Neuroimage 216. https://doi.org/10.1016/j.neuroimage.2020.116797

Kaiser, R.H., Andrews-Hanna, J.R., Wager, T.D., Pizzagalli, D.A., 2015. Large-Scale Network Dysfunction in Major Depressive Disorder: A Meta-analysis of Resting-State Functional Connectivity. JAMA Psychiatry 72, 603–611. https://doi.org/10.1001/jamapsychiatry.2015.0071

Keren, H., Zheng, C., Jangraw, D.C., Chang, K., Vitale, A., Nielson, D., Rutledge, R.B., Pereira, F., Stringaris, A., 2020. Timing matters: The temporal representation of experience in subjective mood reports. bioRxiv 815944. https://doi.org/10.1101/815944

Klimstra, T.A., Kuppens, P., Luyckx, K., Branje, S., Hale, W.W., Oosterwegel, A., Koot, H.M., Meeus, W.H.J., 2016. Daily Dynamics of Adolescent Mood and Identity. J. Res. Adolesc. 26, 459–473. https://doi.org/10.1111/jora.12205

Marco-Pallarés, J., Münte, T.F., Rodríguez-Fornells, A., 2015. The role of high-frequency oscillatory activity in reward processing and learning. Neurosci. Biobehav. Rev. 49, 1–7. https://doi.org/10.1016/j.neubiorev.2014.11.014

Nettle, D., Bateson, M., 2012. The Evolutionary Origins of Mood and Its Disorders. Curr. Biol. 22, R712–R721. https://doi.org/10.1016/j.cub.2012.06.020

Nolte, G., 2003. The magnetic lead field theorem in the quasi-static approximation and its use for magnetoencephalography forward calculation in realistic volume conductors. Phys. Med. Biol. 48, 3637–3652. https://doi.org/10.1088/0031-9155/48/22/002

Nugent, A.C., Ballard, E., Gould, T.D., Park, L.T., Moaddel, R., Brutsche, N.E., Zarate, C.A., 2019a. Ketamine Has Distinct Electrophysiological and Behavioral Effects in Depressed and Healthy Subjects. Mol. Psychiatry 24, 1040–1052. https://doi.org/10.1038/s41380-018-0028-2

Nugent, A.C., Wills, K.E., Gilbert, J.R., Zarate, C.A., 2019b. Synaptic potentiation and rapid antidepressant response to ketamine in treatment-resistant major depression: A replication study. Psychiatry Res. Neuroimaging 283, 64–66. https://doi.org/10.1016/j.pscychresns.2018.09.001

Oostenveld, R., Fries, P., Maris, E., Schoffelen, J.-M., 2011. FieldTrip: Open source software for advanced analysis of MEG, EEG, and invasive electrophysiological data. Comput. Intell. Neurosci. 2011, 156869. https://doi.org/10.1155/2011/156869

Paul, K., Pourtois, G., 2017. Mood congruent tuning of reward expectation in positive mood: evidence from FRN and theta modulations. Soc. Cogn. Affect. Neurosci. 12, 765–774. https://doi.org/10.1093/scan/nsx010

Pierce, J.E., Péron, J., 2020. The basal ganglia and the cerebellum in human emotion. Soc. Cogn. Affect. Neurosci. 15, 599–613. https://doi.org/10.1093/scan/nsaa076

Preuschoff, K., Quartz, S.R., Bossaerts, P., 2008. Human Insula Activation Reflects Risk Prediction Errors As Well As Risk. J. Neurosci. 28, 2745–2752. https://doi.org/10.1523/JNEUROSCI.4286-07.2008

Rutledge, R.B., Skandali, N., Dayan, P., Dolan, R.J., 2014. A computational and neural model of momentary subjective well-being. Proc. Natl. Acad. Sci. 111, 12252–12257. https://doi.org/10.1073/pnas.1407535111

Singer, T., Critchley, H.D., Preuschoff, K., 2009. A common role of insula in feelings, empathy and uncertainty. Trends Cogn. Sci. 13, 334–340. https://doi.org/10.1016/j.tics.2009.05.001

Smith, S.M., Nichols, T.E., 2009. Threshold-free cluster enhancement: addressing problems of smoothing, threshold dependence and localisation in cluster inference. NeuroImage 44, 83–98. https://doi.org/10.1016/j.neuroimage.2008.03.061

Sur, S., Sinha, V.K., 2009. Event-related potential: An overview. Ind. Psychiatry J. 18, 70–73. https://doi.org/10.4103/0972-6748.57865

Tremblay, S., Rogasch, N.C., Premoli, I., Blumberger, D.M., Casarotto, S., Chen, R., Di Lazzaro, V., Farzan, F., Ferrarelli, F., Fitzgerald, P.B., Hui, J., Ilmoniemi, R.J., Kimiskidis, V.K., Kugiumtzis, D., Lioumis, P., Pascual-Leone, A., Pellicciari, M.C., Rajji, T., Thut, G., Zomorrodi, R., Ziemann, U., Daskalakis, Z.J., 2019. Clinical utility and prospective of TMS–EEG. Clin. Neurophysiol. 130, 802–844. https://doi.org/10.1016/j.clinph.2019.01.001

Veen, B.D.V., Drongelen, W.V., Yuchtman, M., Suzuki, A., 1997. Localization of brain electrical activity via linearly constrained minimum variance spatial filtering. IEEE Trans. Biomed. Eng. 44, 867–880. https://doi.org/10.1109/10.623056

Wagner, M.J., Kim, T.H., Savall, J., Schnitzer, M.J., Luo, L., 2017. Cerebellar granule cells encode the expectation of reward. Nature 544, 96–100. https://doi.org/10.1038/nature21726

Woolrich, M., Hunt, L., Groves, A., Barnes, G., 2011. MEG beamforming using Bayesian PCA for adaptive data covariance matrix regularization. NeuroImage 57, 1466–1479. https://doi.org/10.1016/j.neuroimage.2011.04.041

Zrenner, B., Zrenner, C., Gordon, P.C., Belardinelli, P., McDermott, E.J., Soekadar, S.R., Fallgatter, A.J., Ziemann, U., Müller-Dahlhaus, F., 2020. Brain oscillation-synchronized stimulation of the left dorsolateral prefrontal cortex in depression using real-time EEG-triggered TMS. Brain Stimulat. 13, 197–205. https://doi.org/10.1016/j.brs.2019.10.007

