## supplementary material for "Electrophysiological correlates of mood and reward dynamics in human adolescents"

### Exploratory analysis

Exploratory analyses were carried out with a sample of 14 subjects in order to refine hypotheses and analysis methods before pre-registration on OSF (https://osf.io/djw8h).

Hypothesis 1: In the exploratory sample we initially tested whether any frequency band (theta, alpha, beta, beta-gamma, gamma) showed correlation with mood, expectation or reward prediction error. We focused on the 3s “rest” period before the slider scale to rate momentary mood appeared on the screen. From previous literature showing changes in the frontal gamma band associated with depressive symptoms (Fitzgerald and Watson, 2018; Nugent et al., 2019a), we might have expected to see variations in gamma power with mood or reward expectation, but this was not the case in our sample. The clearest significant clusters were observed for the correlation of beta-gamma power (frequency band shown to increase with positive rewards (Marco-Pallarés et al., 2015)) and reward expectation, which we then selected as a testable hypothesis for our confirmatory analysis.

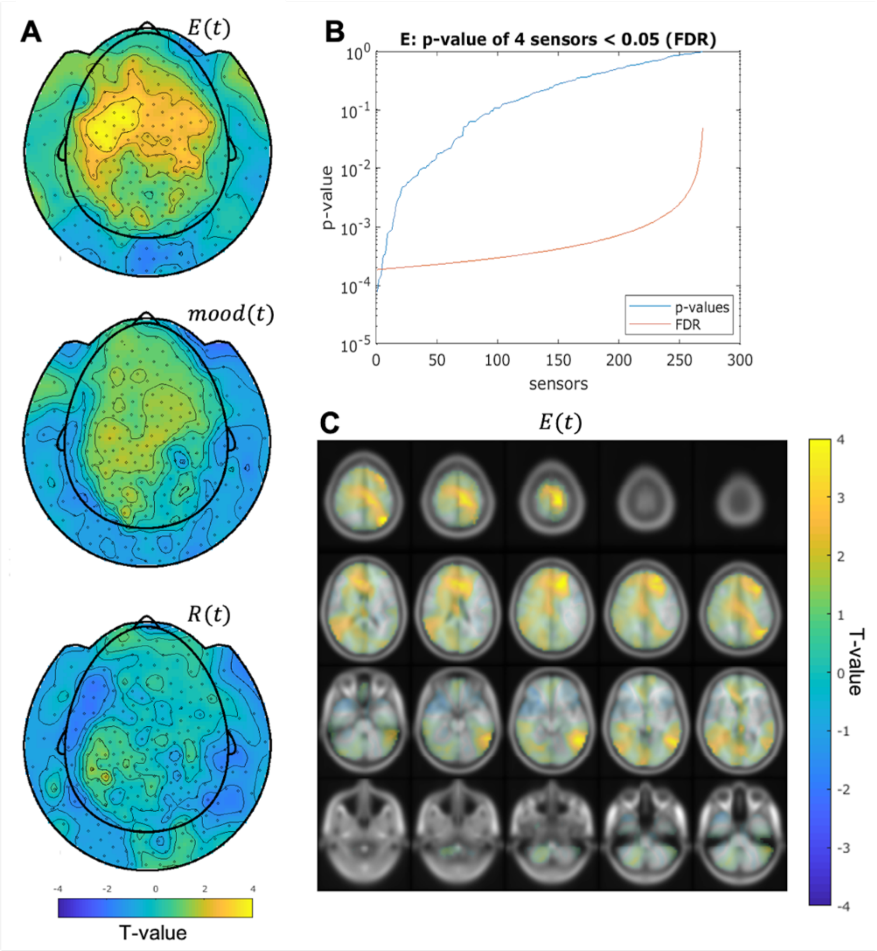


*Figure S1: Results from the exploratory sample (N=14) fitting trial beta-gamma power in the 3s preceding mood-rating to self-reported mood and its primacy model predictors.* ***A.*** *Color maps of t-values of the fixed effects for trial beta-gamma power measured at the MEG sensors. We tested if three separate fixed effects, self-reported mood, expectation and reward prediction error from the primacy model could predict trial level variations in beta-gamma power. Only the expectation predictor $E(t)$ showed significant effects (**B.**), with 4 sensors surviving multiple comparison correction with False Discovery Rate ($\alpha < 0.05, N_{sensors} = 266$, shown in panel).* ***C.*** *Color maps of t-values of the $E(t)$ fixed effect for trial beta-gamma power at the source level. In the exploratory sample we found strong effects in the left superior frontal cortex, ACC, paracentral lobule, left superior and inferior parietal cortex and in the angular gyrus.*

Hypothesis 2: in our exploratory sample we tested if the response to reward feedback was affected by mood, expectation or RPE following the same approach described in the method section for the post-hoc analysis over all 116 ROIs for the AAL atlas. The ROIs showing temporal clusters surviving multiple comparisons (shown in figure S2) were selected for our confirmatory analysis. Unlike previous EEG work, in our exploratory analysis we did not see an effect reconcilable with the Feedback Related Negativity showing correlations with mood or with parameters from our mood primacy model (Paul and Pourtois, 2017).

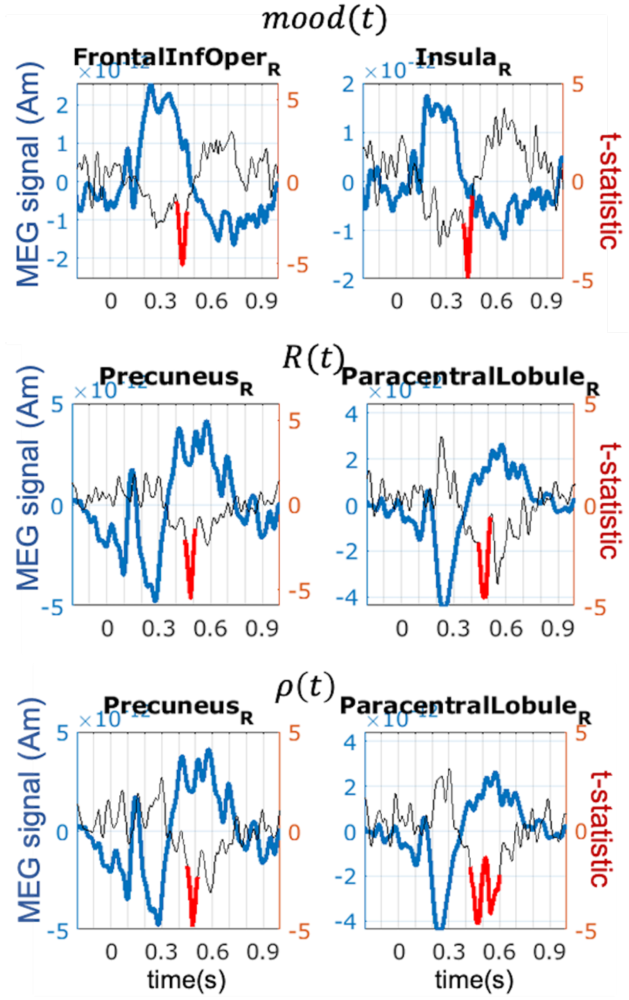


*Figure S2: Response to reward feedback in regions of interest (ROIs) from the AAL atlas from the exploratory sample (N=14). Temporal peaks surviving multiple comparisons over the tested 116 ROIs are highlighted in red.*

Hypothesis 3: We initially selected the 250-400ms time window based on observations on modulations of the P300 component in EEG (De Pascalis et al., 2004; Sur and Sinha, 2009).

In our exploratory sample we tested if the average evoked response 250-400ms following option presentations and choice selection (gamble vs certain amount) showed correlations with mood, choice selection (gamble vs certain amount), or expectation from the primacy model. The only significant effect we observed in our exploratory sample was the reward expectation on the response to gamble option presentation (figure S3) which we then selected as one of our third hypotheses.

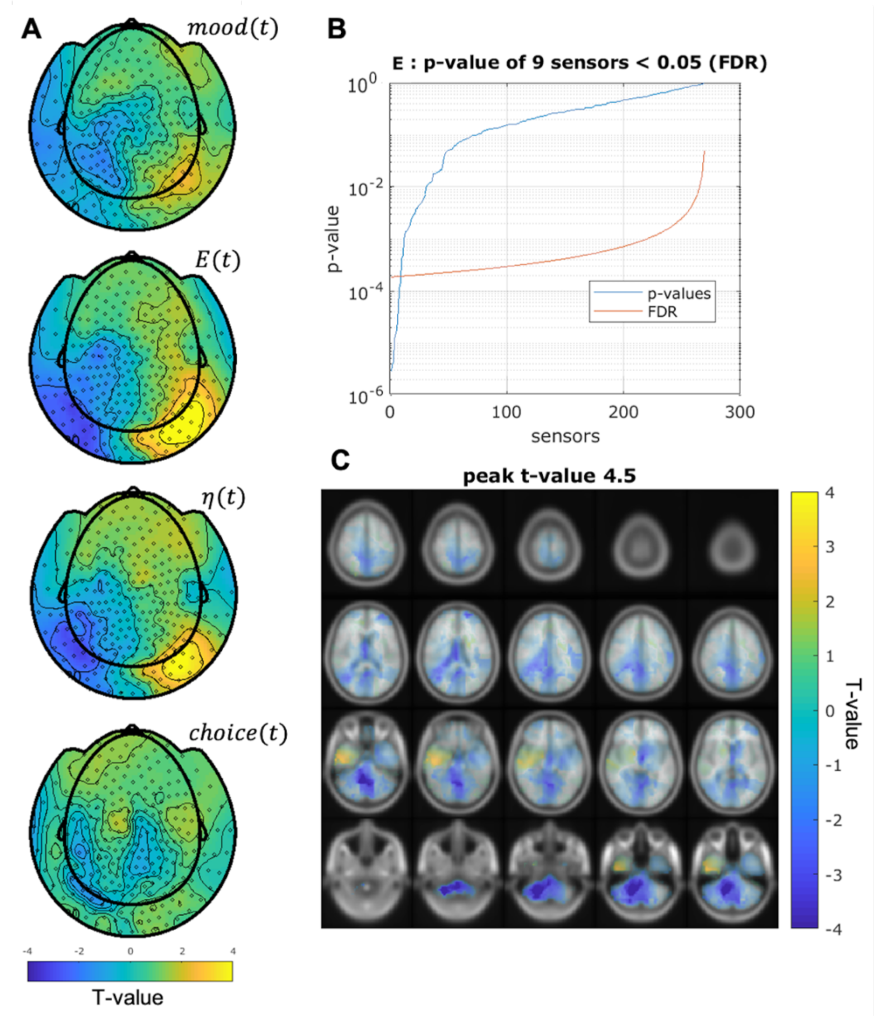


*Figure S3: Results from the exploratory sample (N=14) fitting the MEG signal in the 250-400ms after gambling options presentations to different predictors.* ***A.*** *Color maps of t-values of the fixed effects for MEG signal measured at sensor level. We tested if four separate fixed effects, self-reported mood, expectation, primacy weighted expectation and choice selection could predict trial level variations in beta-gamma power. The expectation predictor* $E(t)$ *and the primacy*

*weighted expectation* $\eta(t)$ *showed significant effects, (**B.**) with* $E(t)$ *having 9 sensors surviving multiple comparison correction with False Discovery Rate (*$\alpha < 0.05, N_{sensors} = 266$*, shown in panel). **C.** Color maps of t-values of the* $E(t)$ *fixed effect for trial beta-gamma power at the source level. In the exploratory sample we found strong effects in the cerebellum.*

**Beta-gamma power correlation with reward expectation: Effect of beamformer regularization and forward model on source localization**

Source localization in MEG (as in EEG) is achieved by reconstructing the magnetic activity detected at the sensors around the subjects the head based on electromagnetic laws describing the magnetic field induced by an electric dipole. The MEG inverse problem is ill-posed as there is not a single possible solution for the electric dipoles inducing a detected magnetic field.

In a beamformer solution one the parameters affecting source localization is the regularization of the data covariance matrix (Brookes et al., 2008). In our hypotheses we selected a high regularization value equal to 5% the maximum singular value of the covariance matrix. Here we explored 2 lower regularization parameters, 1% and 0.2% of the maximum singular value of the regularization matrix to test the stability of our results to variations in the localization technique (Figure S4). We can see that beamformer regularization affects the localization of source clusters where $E_t$ predicts beta-gamma power, with a new cluster in premotor cortex appearing at lower regularization values.

All source localization techniques are also influenced by the physical model we select to calculate the magnetic field outside the head induced by an electric dipole source in the brain. In our hypotheses we selected one of the models most used in the MEG literature, Nolte's single shell approximation (Nolte, 2003). In this section we test another popular MEG forward model, the local spheres model (Huang et al., 1999) (Figure S5). The choice of head model also affects the identification of statistical clusters, with $E_t$ showing overall a lower effect on beta-gamma power with the local spheres model compared to our results with Nolte's single shell approximation. Overall this shows that our multilevel statistical models results at the source level are highly susceptibility to the selection of beamformer parameters. We suggest that for applying a similar analysis technique in future it might be beneficial to apply an adaptive regularization as proposed in (Woolrich et al., 2011) or test localization accuracy at similar SNR level with computational models to determine the best beamformer parameters.

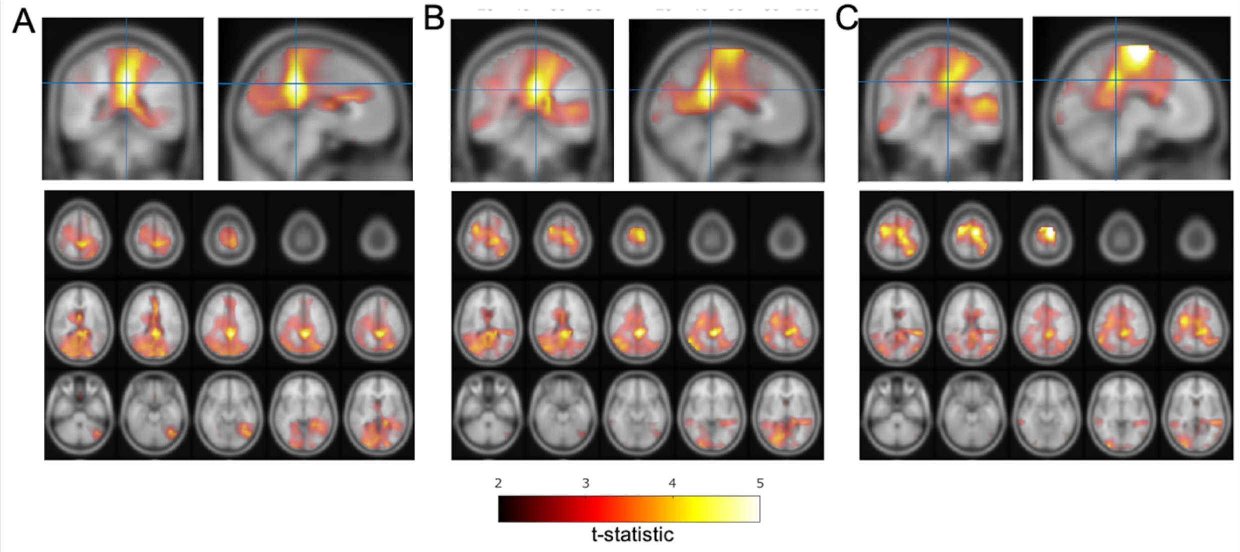


Figure S4: Source space maps showing brain regions where expectation parameters $E_t$ predicts beta-gamma power preceding mood rating. Color bar indicates t-statistic of the fixed effect. A. Source space maps for covariance regularization $\mu = 5\%$ the maximum singular value of the covariance matrix. Top row shows cluster peak in coronal and sagittal views (MNI coordinates [-2 -40 36]mm, left posterior cingulate cortex). The plot on the bottom shows the significant cluster over multiple axial slices.

Plots B. and C. shows the same as A. but for a lower covariance regularization: B. $\mu = 1\%$ and C. $\mu = 0.2\%$ of the maximum singular value of the covariance matrix. Covariance regularization has a large effect on the localization of significant clusters with clusters in left and right premotor cortex appearing at lower regularization values. The cluster in posterior and mid cingulate cortex shifts its peak to [-4 -36 30] for $\mu = 1\%$ (crosshair in top row of B.) and [-8 -36 38]mm for $\mu = 0.2\%$ (top row of C.)

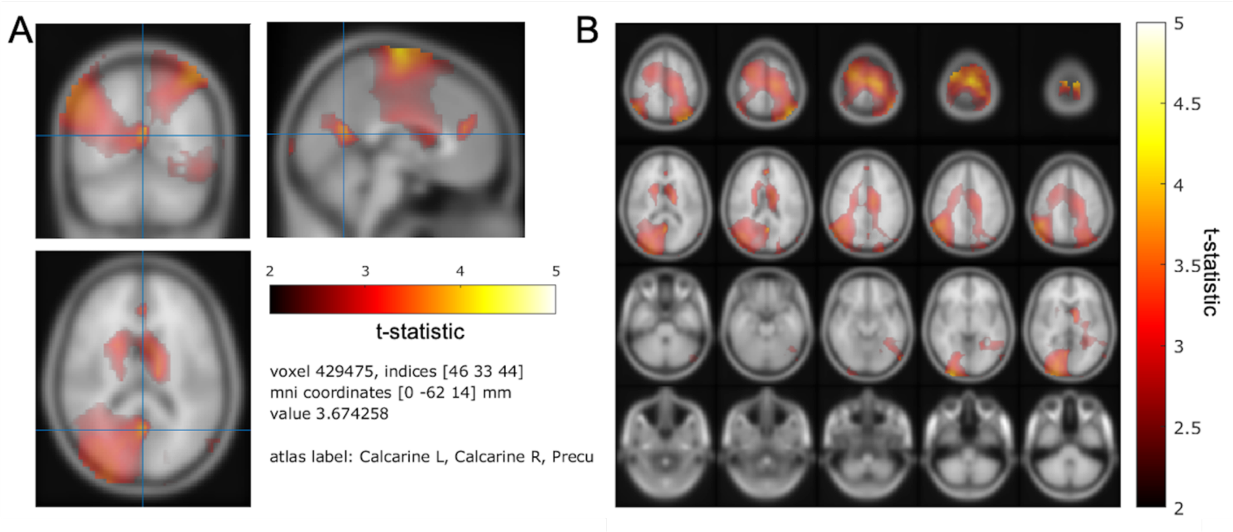


Figure S5: Source space maps from local spheres forward model and beamformer regularization with $\mu = 5\%$ of maximum singular value of the data covariance matrix. Heat maps show brain regions where expectation parameters $E_t$ predicts beta-gamma power preceding mood rating. Color bar indicates t-statistic of the fixed effect.

### Subject Inclusion criteria

#### 1. Youths who meet DSM 5 criteria for Major Depressive Disorder

1.1. Inclusion criteria for Youth with MDD (all must be met):

1.1.1. Ages 11-17 at the time of enrollment in Characterization;

1.1.2. Current diagnosis of DSM-5 Major Depressive Disorder (within the last six months from assessment) which are:

1.1.2.1. Five or more of the following symptoms have been present during the same 2-week period and represent a change from previous functioning; at one of the symptoms is either (1) depressed mood or (2) loss of interest or pleasure.

1.1.2.1.1.1. Depressed mood most of the day, nearly every day, as indicated by either subjective report (e.g., feeling sad, blue, "down in the dumps," or empty) or observation made by others (e.g., appears tearful or about to cry). (In children and adolescents, this may present as an irritable or cranky, rather than sad, mood.)

1.1.2.1.1.2. Markedly diminished interest or pleasure in all, or almost all, activities every day, such as no interest in hobbies, sports, or other things the person used to enjoy doing.

1.1.2.1.1.3. Significant weight loss when not dieting or weight gain (e.g., a change of more than 5 percent of body weight in a month), or decrease or increase in appetite nearly every day.

1.1.2.1.1.4. Insomnia (inability to get to sleep or difficulty staying asleep) or hypersomnia (sleeping too much) nearly every day.

1.1.2.1.1.5. Psychomotor agitation (e.g., restlessness, inability to sit still, pacing, pulling at clothes or clothes) or retardation (e.g., slowed speech, movements, quiet talking) nearly every day.

1.1.2.1.1.6. Fatigue, tiredness, or loss of energy nearly every day (e.g., even the smallest tasks, like dressing or washing, seem difficult to do and take longer than usual).

1.1.2.1.1.7. Feelings of worthlessness or excessive or inappropriate guilt nearly every day (e.g., ruminating over minor past failings).

1.1.2.1.1.8. Diminished ability to think or concentrate, or indecisiveness, nearly every day (e.g. appears easily distracted, complains of memory difficulties).

1.1.2.1.1.9. Recurrent thoughts of death (not just fear of dying), recurrent suicidal ideas without a specific plan, or a suicide attempt or a specific plan for committing suicide

1.1.2.2. Symptoms cause clinically significant distress or impairment in social, occupational/academic, or other important areas of functioning.

1.1.2.3. The episode is not attributable to the physiological effects of a substance or to another medical condition.

1.1.3. Added criteria for Children with MDD entering inpatient treatment. In addition to criteria in 1.1.2 (above), the youth:

1.1.3.1. is failing his/her treatment as defined as a current CGAS score <60.

1.1.3.2. If the child has a psychiatrist, the child's psychiatrist or treater agrees that the child's response to his/her current treatment makes it clinically appropriate to change the child's current treatment.

1.1.3.3. On the basis of record review and interviews with child and parent, the research team agrees that the child's response to his/her current treatment is no more than minimal.

1.1.4. Added criteria for Children with MDD entering outpatient treatment. In addition to criteria in 1.1.2 (above), the youth:

1.1.4.1. Meets criteria for on-going MDD

2. Youths who meet modified DSM criteria for Subthreshold Depression.

2.1. Inclusion criteria for subthreshold depressive disorder are:

2.1.1. Ages 11-17 at the time of enrollment in Characterization;

2.1.2. an episode of depressed mood or loss of interest or pleasure lasting at least 1 week plus

2.1.3. at least two of the seven other DSM-5-associated symptoms for major depression

2.1.4. occurring in the last six months.

3. Inclusion Criteria for Healthy volunteer youths

3.1. Youth 11 to 17 years of age at time of enrollment in Characterization

3.2. Subjects must be competent to assent; parents must be able comprehend and provide permission for their child (consent).

3.3. Youth will be willing to participate in NIMH IRB approved research protocols. Minors will be asked to sign assent forms and their parents will sign the consent form. 3.4. Subjects willing to undergo an evaluation which may include a psychiatric interview, review of medical history (including Tanner staging for minors), pregnancy testing.

3.5. Speaks English

3.6. Has an identified primary care clinician

4. Inclusion Criteria for Parents of enrolled youth:

4.1. Are the biological parent or legal guardian of an enrolled adolescent (who is a healthy volunteer, has s-MDD, or has MDD) participant

4.2. Those of all ages are eligible if they are a parent of a currently enrolled participant c.

**Exclusion criteria (All patients)**

1. Exclusion Criteria for MDD patients (Group 1)

1.1. Meets criteria for schizophrenia, schizophreniform disorder, schizoaffective illness, bipolar disorder, more than mild Autism Spectrum Disorder, Anorexia Nervosa or other severe Eating Disorder.

1.2. Intellectual disability (clinically identified or IQ<70)

1.3. For subjects with major depression or sub-threshold major depressive episode: Symptoms of depression are due to the direct physiological effects of a drug of abuse, or to a general medical or neurological condition by self and parent report.

1.4. Meets DSM-5 criteria for alcohol or substance use disorder (excluding tobacco and nicotine use) within the last three months. This is determined solely by clinical interview of child and parent (e.g. KSADS).

1.5. Current active suicidal ideation (i.e., presence of intent for engaging in suicidal behaviors). Youths with passive suicidal ideation and/or past active suicidal ideation are still eligible.

1.6. Participants with repeated self-harm occurring in the context of inter-personal conflict.

1.7. NIMH IRP Employees/staff and immediate family members will be excluded from the study per NIMH policy.

2. Exclusion criteria for youths meeting modified DSM criteria for Subthreshold Depression (Group 2):

2.1. Intellectual disability (clinically identified or IQ<70).

2.2. Any serious medical condition (such as epilepsy, heart disease requiring medication) by self and parent report.

2.3. Past or current diagnosis of a manic or hypomanic episode, major depression), schizophrenia, schizophreniform disorder, schizoaffective illness, Tourette Disorder, or Autism Spectrum Disorder, Anorexia Nervosa or other severe Eating Disorder.

2.4. Meets DSM-5 criteria for alcohol or substance abuse within the last three months by self and parent report.

2.5. NIMH IRP Employees/staff and immediate family members will be excluded from the study per NIMH policy.

3. Healthy volunteer youth exclusion criteria:

3.1. Intellectual disability (clinically identified or IQ<70).

3.2. Any serious medical condition (such as epilepsy, heart disease requiring medication) by self and parent report.

3.3. Past or current diagnosis of any mood disorder (manic or hypomanic episode, major depression), anxiety disorder (except specific phobia), Obsessive Compulsive Disorder (OCD), Post-Traumatic Stress Disorders (PTSD), Conduct Disorder, schizophrenia, schizophreniform disorder, schizoaffective illness, Tourette Disorder, or Autism Spectrum Disorder.

3.4. Meets criteria for subthreshold depression (as defined above)

3.5. Meets DSM-5 criteria for alcohol or substance abuse within the last three months by self and parent report.

3.6. NIMH IRP Employees/staff and immediate family members will be excluded from the study per NIMH policy.

4. Parents of enrolled participants exclusion criteria:

4.1. Parents who are unable to understand or read English well enough to complete the study interview and tests.
4.2. Parents who are a current NIMH employee, or staff member, or a family member of an NIMH employee. NIMH IRP Employees/staff and immediate family members will be excluded from the study per NIMH policy.

Patients will be recruited locally and nationwide through mailings to selected physicians, announcements in newsletters, and contacts with support groups and approved websites. Color or black and white copies of ads and flyers may be used based on publication and cost. Travel expenses for the patient and accompanying parent will be reimbursed. Healthy volunteers for the control group will be recruited through advertisements in the local media and websites. Control subjects will be compensated according to NIH guidelines.

Our recruitment program seeks to increase the awareness of our studies among a broad demographic (e.g. patients and families, scientific community, mental health clinicians) through a range of recruitment materials and venues.

**Research and Travel Compensation**

All volunteers will be compensated for time and research-related inconveniences. Compensation will be prorated for parts completed if subjects do not complete the entire study. Payment will be provided in the name of the child.

Lodging and travel compensation for those living within the United States and over 50 miles from NIH will be provided. Reimbursement for travel expenses will be provided in accordance with NIH guidelines. Local taxi fare will also be reimbursed, when needed. An escort fee also is provided.
